# A neuron that regulates locomotion makes a potential sensory cilium lying over the *C. elegans* egg-laying apparatus

**DOI:** 10.1101/2022.09.19.508547

**Authors:** Nakeirah T.M. Christie, Michael R. Koelle

## Abstract

The neural circuit for *C. elegans* egg laying has been studied intensively for decades, yet it is not clear that its known components can account for how egg-laying and locomotion behaviors are coordinated. We found that the two PVP neurons, which release neuropeptides that promote roaming locomotion, make previously-undescribed branches that terminate in large wing-shaped endings directly over the egg-laying apparatus. The PVP branches occur in hermaphrodites but not males and develop during the L4 larval stage when the egg-laying system also develops. The PVP wing is located at the junction between the uterus and the vulva, adjacent to neurons that control egg laying, and surrounded by cells that we found label with a glial marker. The morphology of the PVP wing and its envelopment within possible glial cells are consistent with the hypothesis that the PVP wing is a sensory cilium. Although PVP is reported to express sensory receptor homologs, we have been unable to detect PVP expression of more specific markers of neural cilia, and we have also not detected strong PVP defects in the *daf-19* mutant, which does show defects in known neural cilia. The PVPs are extraordinarily sensitive to expression of transgenes, which cause developmental and possibly functional defects in these neurons. This has prevented us from recording or manipulating PVP activity to determine its functional roles. Thus, the intriguing hypothesis that PVP is a sensory neuron that might coordinate egg laying and locomotion will remain speculative until better methods to manipulate PVP can be developed.

## Introduction

Neural circuits are functional units in the brain consisting of groups of neurons that respond to stimuli and signal to each other to produce dynamic patterns of activity (Bargmann and Marder 2013). Defects at the level of neural circuit function are thought to result in a variety of highly prevalent mental health problems. For example, it is estimated that over a fifth of U.S. adults will experience a mood disorder at some point in their life (Kessler *et al*. 2005). It remains difficult to understand how abnormalities in neural circuits arise and result in such disorders because the mechanisms that underlie neural circuit function remain poorly understood. Currently there is no neural circuit in any organism in which it is understood how the complete set of signals among the all the neurons involved produces the dynamic pattern of activity that carries out the function of the circuit.

One approach to advance our understanding of neural circuits is to intensively study small neural circuits of genetically-tractable invertebrates, with the hope that deeply understanding a few such model circuits will reveal general principals by which all neural circuits operate. With only 302 neurons, the nematode *C. elegans* offers the opportunity to completely describe all the components of individual neural circuits. Alongside a physical connectome which traces each neuron’s processes and the ∼7000 physical synapses they make (Albertson and Thompson 1976; White *et al*. 1986; Xu *et al*. 2013), there is also a detailed map of which neurons release what neurotransmitter (Serrano-Saiz *et al*. 2013; Pereira *et al*. 2015; Gendrel *et al*. 2016).

Within *C. elegans*, one of the best-characterized model neural circuits is the egg-laying circuit. Not only does it have only ∼12 neurons and 16 muscle cells, but it also has a very simple output (i.e. an egg being laid) ideal for quantitative behavioral analysis (Schafer 2005). Despite the fact that each neuron known to be part of the circuit has had its activities and functions investigated in detail (Collins and Koelle 2013; Collins *et al*. 2016; Brewer *et al*. 2019; Kopchock *et al*. 2021), there remain behaviors produced by this circuit that cannot be explained by these neurons. For instance, previous research shows that the presence of eggs in the uterus stimulates activity of the egg laying circuit, but no mechanism for the circuit to sense these eggs is known (Collins *et al*. 2016). Eggs tend to be laid at a particular phase of locomotor body bends (Collins *et al*. 2016), and animals do not lay eggs when immobilized, but the neural mechanisms responsible coordinating egg laying locomotion behavior remain unknown. Additionally, a variety of factors in the environment (e.g. the presence of food) appear to regulate egg laying (Dong *et al*. 2000), but the mechanisms used to sense these factors and relay their presence to the egg-laying circuit remain unclear. These unexplained phenomena imply that there are likely additional cells that regulate activity of the currently-recognized components of the egg-laying circuit. As long as these additional components of the egg-laying system remain unknown, it will not be possible to obtain a full mechanistic understanding of how activity of the egg-laying circuit is controlled, limiting the utility of the egg-laying circuit as a model for understanding neural circuit function and dysfunction in general. In this work, we have characterized a pair of PVP neurons that may constitute a new component of the *C. elegans* egg-laying system.

## Materials and Methods

### Strains and culture

The supplemental material contains a list of *C. elegans* strains used in this study. *C. elegans* strains were cultured at 20°C on NGM agar plates with *E. coli* strain OP50 as a nutrient source (Brenner 1974). Genetic crosses were by standard methods (Fay 2013). Extrachromosomal-array transgenic strains were generated through standard microinjection (Evans 2006). Chromosomal integration of such transgenes was performed via UV/TMP mutagenesis (Fernandez *et al*. 2020), and integrated strains were backcrossed to N2 to remove background mutations as indicated in Table 1.

### Transgenes

The supplemental material describes all transgenes used in this study. A GFP reporter plasmid, pEXP294, expressed exclusively in PVP within L4 and adult animals was a gift from Richard Ikegami. In this plasmid, the promoter used (*PVPp*) is derived from 699 bp of the *C. elegans* dystroglycan (*dgn-1*) gene, (primers F:catggggatccggacaatgagaatgag; R:gcctttttgtcttatatacattttttttagaccgttcaca). To increase expression of GFP the Syn21 translational enhancer and AcNPV p10 3’UTR (Pfeiffer *et al*. 2012). The *PVPp::gfp* transgene used to visualize PVP also co-expressed a histamine-gated chloride channel in PVP, although histamine was not applied to the animals to activate this channel in any experiments shown in this article. To express other proteins in PVP, we cloned the *dgn-1* promoter and AcNPV p10 3’UTR into the vector pPD49.26 (from the Andrew Fire plasmid kit, Addgene Kit #1000000001) to generate plasmid pNTC2, wherein we were able to drop coding sequences of interest into the multiple cloning site downstream of the promoter along with the Syn21 translational enhancer to express proteins of interest in PVP (Table 2).

### Confocal Imaging

Animals were mounted on agarose pads on microscope slides Superfrost (Thermo Fisher Scientific) and a 22×22-1 microscope cover glass (Fisher scientific). The 2% agarose pads contained containing 25 mM of sodium azide (Sigma Aldrich) and 120mM of 60% w/v iodixanol (Optiprep) (Sigma Millipore) to reduce refractive index mismatch (Boothe *et al*. 2017). Images were acquired on an LSM 880 using a 40*x* water immersion objective (Carl Zeiss), with either standard confocal imaging, spectral imaging (lambda stacking) to filter out autofluorescence, or fast Airyscan SR mode, as indicated.

### Developmental Time Course

Animals carrying transgenes that label PVP were picked on a dissecting microscope at various developmental points within the L4 larval stage and mounted for confocal imaging, recording both fluorescent and brightfield images. Using the brightfield image, animals were sub-staged according to vulval morphology (Mok *et al*. 2015), binning L4.0-L4.2 as “early L4”, L4.3-L4.5 as “mid-L4”, and L4.6-L4.9 as “late L4.”

## Data Availability

Strains and plasmids are available upon request. The supplemental material contains detailed descriptions of all these reagents.

## Results

### The PVP neurons produce previously-uncharacterized branches at the midbody

While examining the *C. elegans* egg-laying system in a set of 20 strains in which the Green Fluorescent Protein (GFP) labels various sparse subsets of neurons (Fernandez *et al*. 2020), we noticed a neural structure that had not been previously described in the published literature. In two of the strains, a neural branch extended from the ventral nerve cord at the midbody near the vulva and terminated in a striking wing-shaped neural projection. Each of these two strains showed GFP labeling of its own set of ∼20 types of neurons, with the overlap between these two sets of being a pair of neurons with their cell bodies in the tail at positions consistent with those of the left and right PVP neurons. Using a transgene (PVPp::GFP) that drives GFP expression exclusively in PVP from a fragment of the dystroglycan promoter, we confirmed that the midbody neural branch structure was indeed produced by the PVP neurons (Figure 1A-B). The PVP neurons were previously classified based on reconstruction from electron micrograph (EM) serial sections as an interneuron pair with cell bodies located in the pre-anal ganglion that send processes anteriorly along the left and right ventral nerve cord (VNC) before terminating in the nerve ring located within the head of the animal (White *et al*. 1986). The midbody branches we observed, one produced by each PVP, extended dorsally from the VNC near the vulva and terminated over the uterus in a wing-like structure (Figure 1A.) The branches were seen in hermaphrodites but not found in males (Figure 1B-C).

**Figure 1.**
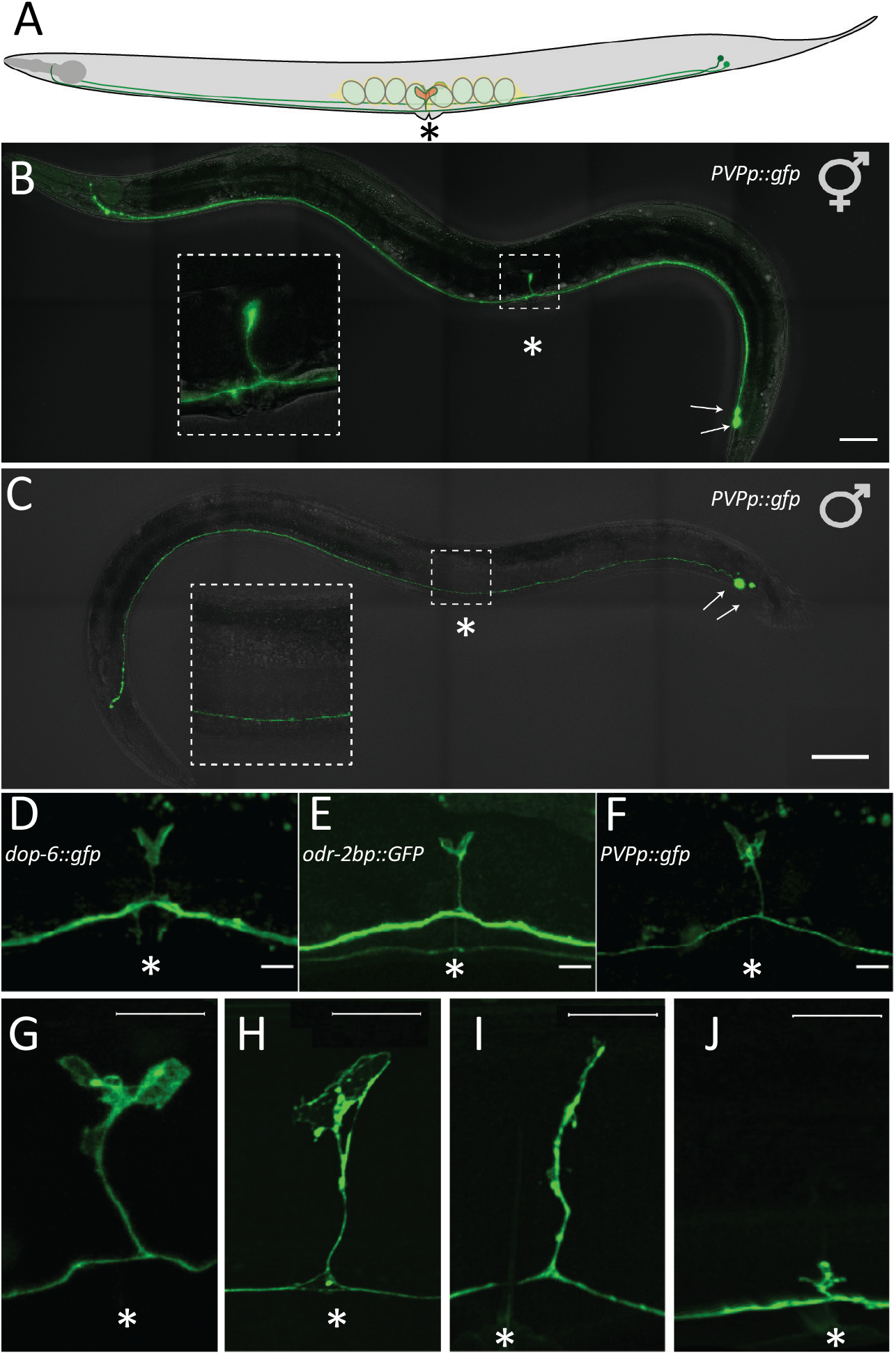
The PVP neuron pair produce a previously uncharacterized branch with wing-shaped endings at the midbody of adult hermaphrodite *C. elegans*. (A) Diagram of PVP neurons within a *C. elegans* adult hermaphrodite. The left (light green) and right (dark green) PVPs have cell bodies near the tail (dots), and processes (lines) that cross as they go anteriorly along the ventral nerve cord to ultimately terminate in the head. Each PVP makes a branch near the vulva that terminates in a wing-like structure (red). In this and subsequent figures, an asterisk is placed just ventral to the vulval opening as a reference point. (B) PVP neurons visualized in an adult hermaphrodite carrying a transgene that expresses GFP from a PVP specific promoter (*PVPp*). Arrows indicate the PVP cell bodies. Inset magnifies the vulval region, where one PVP branch and wing are visible in the focal plane shown. Scale bar, 50 μm. (C) Adult male with the male equivalents of the PVPs visualized as in (B). Inset magnifies the midbody region. The male neurons have no branch or wing. (D-F) Vulval region of adult hermaphrodites expressing GFP in PVP from three different promoters. PVP branches and wings appear similar in each. Scale bars, 10 μm. In (D) an (E) processes from neurons other than PVP are visible in the images. (G-J) Airyscan (super resolution) images of PVP in the vulval region demonstrating morphological variations in the branch and wing. Scale bars, 10 μm.

The PVP midbody branches were not described in the original EM reconstruction of the *C. elegans* nervous system (White *et al*. 1986), which caused us to be concerned that the GPF-labeled PVP branch we saw could be an artifact of the PVPp::GFP transgene. Significant deviations from the results of White *et al*. (1986) are almost unheard of, and these results have rather been confirmed by a vast body of subsequent work (e.g. Witlievet *et al*. 2021). Therefore, we obtained a set of three promoters that are expressed in PVP (among other cells) and used them to express GFP and/or a red fluorescent protein in transgenic animals. Figure 1D-F show representative images from animals expressing GFP from three different promoters, in each cases showing a similar branch. We noted that while the PVP branches generally ended in ‘winged’ shapes, we saw a wide variety of morphologies and lengths in different individual animals (Figure 1G-J).

### PVP morphology and function are extremely sensitive to transgene expression

Originally, we intended to study the function of PVP and its branch by transgenically expressing a variety of proteins in PVP to record PVP activity (e.g. GCaMP) or manipulate PVP activity (e.g. channelrhodopsin or a histamine-gated chloride channel; HisCl), similar to studies our laboratory has carried out of other neurons of the egg-laying system (e.g. Collins *et al*. 2016). However, in the course of attempting to execute these experiments, it became evident that the PVPs are extremely sensitive to these transgenes. It was difficult to generate viable lines driving expression of any protein in PVP besides GFP. Expression of red fluorescent proteins, such as mCherry and TagRFP, typically led to malformed PVP cells that often showed punctate red fluorescence, features that might indicate defects in PVP development or degeneration of the PVPs. Figure 5G shows the only image presented in this work of a red fluorescent PVP neuron, and this is one of the healthiest-looking red fluorescent PVPs we have seen.

Of the transgenic strains mentioned above that we attempted to generate for functional studies assays of PVP, only one proved viable: a strain carrying a transgene that expressed both GFP and the HisCl channel from the PVP-specific promoter. This transgene was chromosomally integrated and outcrossed to the wild type to produce a clean genetic background, resulting in the “PVPp::HisCl/PVPp::GFP” strain. The GPF labeling of PVP in this strain is shown in Figure 1, and the PVP cells appear relatively healthy in this strain.

Transgenic worms expressing the HisCl channel can be treated with histamine to inactivate neurons that express the channel (Pokala *et al*. 2014). We assayed the PVPp::HisCl/PVPp::GFP strain for changes in behavior after treatment with histamine. We found that egg-laying behavior was relatively normal with or without histamine treatment, but that a locomotion behavior termed “roaming” (Flavell *et al*. 2013) appeared to be defective regardless of whether animals were treated with histamine or not (data not shown). Flavell *et al*. (2013) had previously shown that PVP neurons release the PDF-1 neuropeptide to promote roaming, and our results suggest that the PVPp::HisCl/PVPp::GFP transgene inactivated this function of PVP neurons even without the need to open the HisCl channels with histamine. Thus, we conclude that likely all of our transgenes that are expressed in PVP, even those that show the most healthy-appearing PVP morphology, in fact compromise the function of PVP.

The sensitivity of PVP function to transgenes raised the question of whether the morphologies of the branch and wing structures of PVP were also affected by transgenes used to visualize them. While we originally observed the PVP wing-structure extending dorsally from the VNC at the midbody (Figures 1B, 1D-F, and 2A), we also sometimes observed PVP wing-structures curved ventrally towards the vulva (Figure 2B). This occurred a little less than 50% of the time in PVPp::HisCl/PVPp::GFP animals (9 out of 20 animals examined; Figure 2C). We re-examined other transgenic lines with GPF expression in PVP for this phenomenon (Figure 2D-Figure 2F). In the *odr-2p*::GFP animals, we observed dorsally-oriented PVP wing structures 54% of the time, ventral orientations 26%, and no visible branches for the remaining 20% (n=15). We also expressed GFP under a the *ocr-3b* promoter, which like our original *dgn-1*-derived promoter, is very specific to PVP in adult animals (Lorenzo *et al*. 2020). These *ocr-3p*::GFP animals had fluorescent PVP neurons in which no branch or wing structure was visible (n=15), and on at least one occasion, the main ventral cord process of the PVP neurons terminated at a point between the vulva and the head and failed to reach its normal more anterior destination in the nerve ring of the head (Figure 2F). All this together suggests the PVP branch and wing structures are sensitive to transgene expression, and that the variations in branch and wing in animals that carry the same PVPp::HisCl/PVPp::GFP transgene (Figures 1G-1J) may be due to effects of the transgene itself. That being the case, the most common PVP morphology seen, and the one reproduced in multiple independent transgenic lines, is that with a dorsal branch terminating in a large wing-like structure, as seen in Figures 1D-1G. This most likely represents the morphology of a healthy PVP.

**Figure 2.**
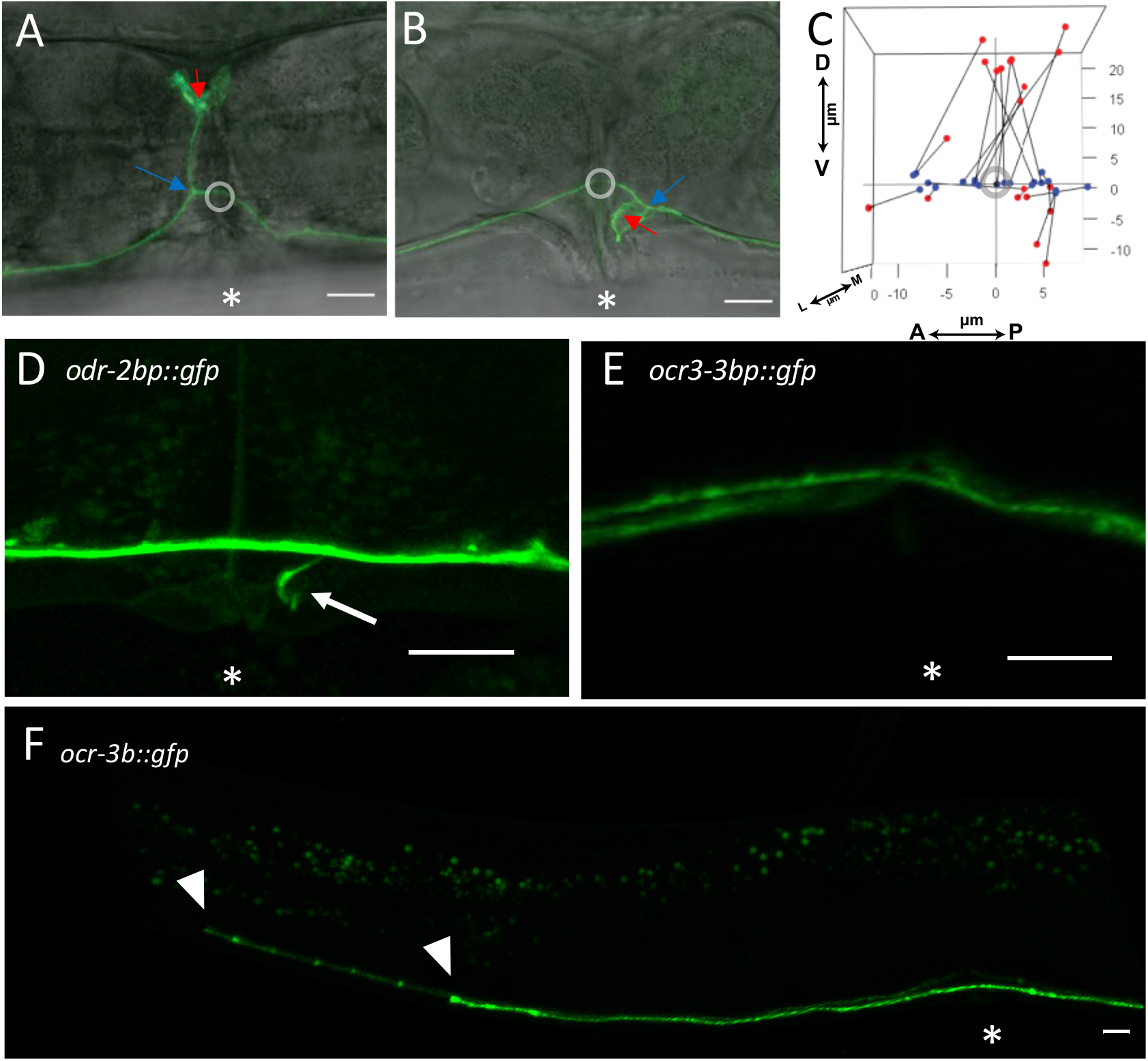
*P*VP neuron*s* are sensitive to transgene expression. Representative confocal images of a dorsally-extending PVP branch/wing structure (A) and a ventrally-oriented PVP branch/wing structure (B). Grey circles enclose the points where main PVP process crosses the vulval slit, blue arrows indicate the PVP branch point, and red arrows indicating the approximate center of the PVP wing structure. (C) Graph showing the 3-dimensional positions of the branch points (blue dots) and wing centers (red dots) for 20 PVP neurons, with these two reference points for each PVP neuron connected by a black line. Axes of the graph are the <u>D</u>orsal/<u>V</u>entral, <u>A</u>nterior/<u>P</u>osterior, and <u>L</u>ateral/<u>M</u>edial axes of the animal, and the origin of the graph (grey circle) is the point at which the main PVP process crosses the vulval slit. (D-F) PVPs transgenically expressing GFP display morphological variations that can depend on the promoter (indicated on each panel) used to express GFP. (D) A ventrally-directed PVP branch (arrow); (E) PVP main processes pass the vulval region without making branches; (F) PVP main processes with no branch at the vulval region that terminate (arrowheads) before reaching their normal destinations in the head. Scale bars, 10 μm in all panels.

### The PVP branch and wing develops at the L4 stage and lie within the egg-laying system

Because the PVP branch and wing lies near the egg-laying system of adult hermaphrodites, we asked if this structure develops at the L4 larval stage, when the cells of the egg-laying system differentiate (Desai *et al*. 1988). We imaged animals throughout the L4 larval stage (Figure 3), visualizing PVP with the PVPp::HisCl/PVPp::GFP transgene, and visualizing the outlines of vulval and uterine cells that form the main structural elements of the egg-laying system with an *ajm-1:mCherry* transgene that labels the apical junctions of these cells (Köppen *et al*. 2001). We staged the animals according to vulval morphology as early, mid, or late L4 (Mok *et al*. 2015). Early L4 animals showed no PVP branches (n=10), about 40% of the animals had some sort of branch in mid L4 (n=10), and all animals in the late L4 stage showed branches (n=10).

**Figure 3.**
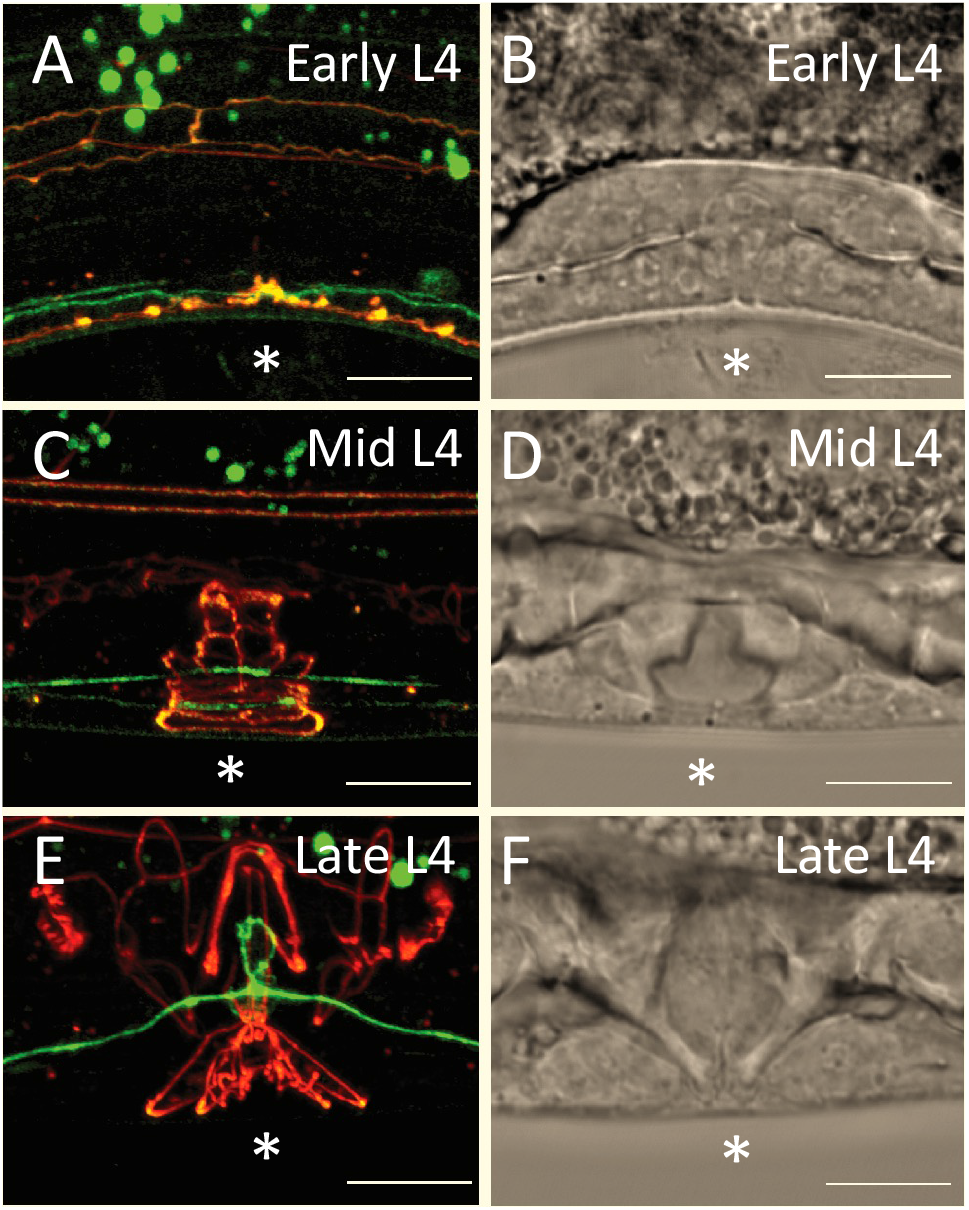
Time course of PVP development within the L4 larval stage. Region of the developing vulval in L4 animals carrying the *PVPp::gfp* transgene to visualize PVP neurons and the *ajm-1::mCherry* transgene to label the apical junctions of epithelial cells, imaged by confocal fluorescence. Left panels show fluorescence images that reveal the developing morphology of the PVP (green) and the developing morphology of structural cells of the vulva and uterus (red). Right panels show brightfield images of the same animals, in which morphology of the developing vulva is also seen. (A-D) In early L4 and mid L4 animals the PVP passes by the vulva without consistently showing branches. (E-F) Late L4 animals have consistently initiated PVP branching. Scale bars, 10μm.

We examined the anatomical relationship of the PVP branch and wing to cells of the egg-laying system in adult animals using GFP to mark PVP and mCherry to mark specific cells of the egg-laying system. As noted previously, some variations in PVP morphology may be artifacts of the transgenes used to visualize PVP, so the most constant features of PVP anatomy are likely the most meaningful. The point at which the branch leaves the main ventral cord process at points that may be either just anterior or just posterior to the vulval opening, ranging over a ∼15 μm span relative to the vulva (Figure 2C). Despite this variation, the wing structure in which the branch terminates is consistently located at the junction between the vulva and the uterus. Figure 4A schematizes the structural cells of the adult egg-laying system whose apical junctions are labeled by the *ajm-1::mCherry* transgene. Unlaid eggs are stored in the uterus, which is a tube constructed from a set of uterine toroid (ut) cells and a uterine seam cell. Eggs are laid by passing through connection between the uterus and vulva formed by the dorsal uterine (du) cell and the uterine-vulval (uv) cells. Finally, the eggs proceed out of the animal through a tube formed by the vulval toroid (vt) cells. Despite variations in origin and path of the PVP branch, when PVP branches proceed dorsally they consistently (10/10 animals examined) terminate in wings that lie over junction of the uv and du cells (Figure 4B).

**Figure 4.**
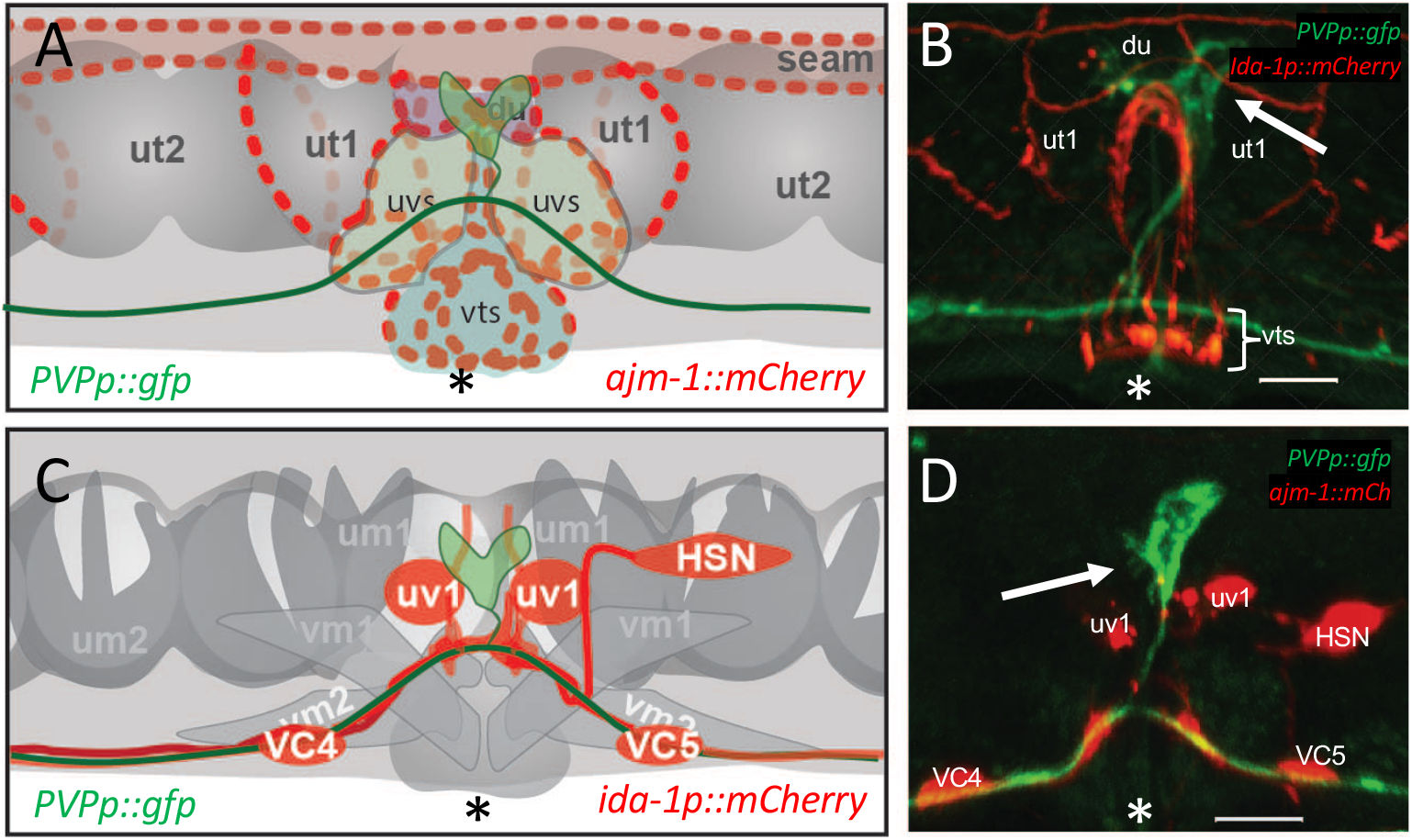
PVP branches and wings lie at the connection between the vulva and uterus of the egg-laying system. (A-B) Schematic (A) and confocal image (B) of on the left side of the egg-laying system of an animal expressing *PVPp::gfp* (green) and *ajm-1::mCherry* (red). Cells outlined by the *ajm-1::mCherry* marker include the uterine toroid cells (ut1 and ut2), vulval toroid cells (vts), dorsal uterine cell (du), and uterine seam cell (seam). The PVP wing (arrow in B) consistently lies over the du cell. (C-D) Schematic (C) and confocal image (D) of the left side of the egg-laying system of an animal expressing *PVPp::gfp* (green) and *ida-1p::mCherry* (red). Schematic indicates muscle cells and neurons of the egg-laying system, including the uterine muscles cells (um1 and um2), vulval muscle cells (vm1 and vm2), uv1 neuroendocrine cells, and VC and HSN neurons. *ida-1p::mCherry* labels the neural cells. PVP wings (arrow) are centrally located between and dorsal to the uv1s. Scale bars, 10 μm.

**Figure 5.**
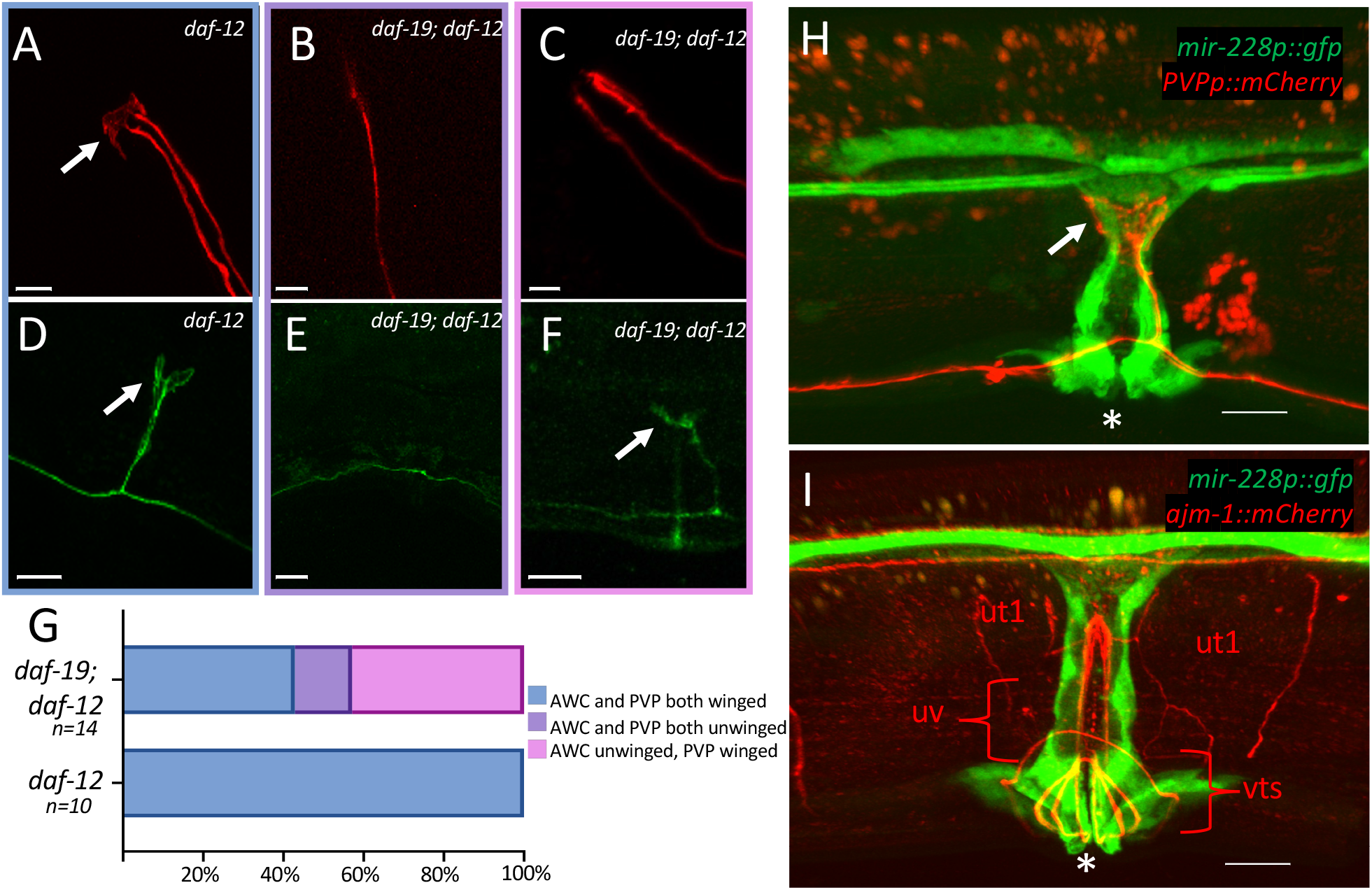
PVP branches and wings are only rarely affected by a mutation that disrupts sensory cilia, but lie among cells that express a glial marker. (A-F) The *PVPp::gfp* transgene that labels the PVP (green) and an *odr-1p::mCherry* transgene that labels the AWC and AWB ciliated sensory neurons (red) were both crossed into the *daf-12* mutant background (A, D), in which cilia are normal, and into the *daf-19; daf-12* double-mutant background (B, C, E, F), in which sensory cilia are disrupted. In *daf-12* animals, the AWC sensory dendrites terminate in winged cilia (A, arrow), and the PVP branch terminates in a similar-appearing winged structure (D, arrow). In the *daf-19; daf-12* double-mutant, almost 60% of AWC neurons lack visible winged cilia (B, C). In this same background, PVP branches and wings were missing in only 2/14 animals examined (E), and the remaining animals showed PVP branches and wings (F, arrow indicates wing). (G) Quantification. (H) The PVP branch and wing (red) lie within cells labeled by the *mir-228p::gfp* glial marker (green). Scale bar, 10 μm. (I) The *ajm-1::mCherry* marker (red) serves as a reference to help identify the cells expressing *mir-228p::gfp* (green). Scale bar, 10 μm.

Figure 4C schematizes neurons of the egg-laying system that are labeled by a IDA-1::mCherry transgene. The uv1s are neuroendocrine cells that form part of the uterine-vulval connection. As shown in Figure 4B, PVP wing structures consistently lie just dorsal to the uv1s and between the anterior and posterior uv1 cells. This again places the uv1 wings directly over the uterine-vulval connection. On occasion we have observed smaller tendrils or branches extending off of the main wing structure extending towards the HSN and uv1 neurons, close to the large synaptic junctions made by the HSN.

### The PVP wing structures show some but not all features of sensory cilia

The anatomy of the PVP branch and wing are reminiscent of the anatomy of dendrites and ciliated endings of sensory neurons in *C. elegans*. In particular, the PVP wing appears similar to the wing-like ciliated ending AWC amphid neuron of the head. We wondered if the PVP wing structures could be cilia as well. We introduced both the *PVPp::gfp* transgene and an *odr-1p::mCherry* transgene, which labels the AWC and AWB sensory neurons, into a genetic background (*daf-19; daf-12*) that disrupts the structures of sensory cilia (Senti and Swoboda 2008). The *daf-12* mutation disrupts cilia, while the *daf-12* mutation is necessary to suppress otherwise lethal effects of the *daf-19* mutation. Thus, the *daf-12* single-mutant background is used as a control in this experiment. In this *daf-12* control background, the AWC and PVP wing-like endings appeared similar and were present in both cell types in all animals examined (Figures 5A, 5D, and 5G). In the *daf-19; daf-12* “cillialess” background, wing-like cilia were absent from almost 60% of AWC neurons examined, although the sensory dendrites were still present. However, in these same *daf-19; daf-12* animals, the wing structure was still present in 12/14 of PVP neurons examined.

We examined a variety of *C. elegans* strains available that were reported to mark neural sensory cilia in young larval stages with fluorescent proteins, but these strains either did not show visible fluorescence in PVP or did not visibly express the fluorescent reporter protein in any neurons at the L4 or adult stages (see File S1). This line of experimentation thus failed to provide any further support for the hypothesis that the PVP wings are sensory cilia.

Sensory dendrites and cilia in *C. elegans* are surrounded by glial cells (Shaham 2006), and we tested whether the PVP dendrite and wing share this feature. We crossed a *PVPp*::*mCherry* transgene that labels PVP with red fluorescence into a strain carrying a transgene (*mir-228p::gfp*) that labels all glial cells with green fluorescence (Fung *et al*. 2020; Pierce *et al*. 2008). In the resulting double-labeled strain, we saw that the PVP branch and wing were surrounded by cells that expressed the glial marker (Figure 5H). Previously, it has been noted that the *mir-228p::gfp* marker labels cells near the vulva (Pierce *et al*. 2008, Fung *et al*. 2020), but these cells were not identified and had not been considered to be candidates to be *bona fide* glial cells since no neural sensory endings at the vulva were known. We crossed the *mir-228p::gfp* transgene to the *ajm-1::mCherry* transgene that outlines structural cells of the egg-laying system (Figure 5I). While a definitive identification of the midbody cells that express the *mir-228* glial marker will require a more detailed analysis, it appears that they include vulval toroid (vt) cells, one or more of the uterine-vulval (uv) cell types, as well as additional uterine cells. Regardless of the exact identity of these cells, the fact that the PVP branch and wing are surrounded by cells that express the *mir-228* glial marker supports the hypothesis that the PVP branch is a sensory dendrite and the PVP wing is a sensory cilium.

## Discussion

The goal of this work was to investigate previously uncharacterized neural branches near the *C. elegans* egg-laying system. The neurons that generate egg-laying behavior constitute an intensively-studied model neural circuit, so the discovery of a new component of this circuit could enhance the power of this model system to provide insights into how neural circuits in general function. It has been a great rarity to discover new features of any C. *elegans* neuron since the ground-breaking study of White et al. (1986) provided a virtually complete description of the anatomy and connectivity of all neurons in the hermaphrodite worm. However, it first was observed in 1999 that the PVP neurons made previously undescribed branches near the hermaphrodite vulva (O. Hobert and D. Hall, personal communication). We determined that neural branches we observed near the hermaphrodite vulva indeed arise from the PVP neurons, and we present here our initial characterization of the structure of the PVP branches and their possible functions.

We found that the PVP neurons have hermaphrodite-specific branches whose development and structure suggest they function in some aspect of reproduction. The PVP neurons were previously described as being born near the tail during embryonic development and then extending processes to the head, with the right PVP (PVPR) process crossing the ventral midline to serve as a pioneer for the left tract of the ventral nerve cord (Durbin, 1987; Wadsworth and Hedgecock, 1996; Wadsworth et al. 1996). The PVPs were also previously known to be sexually dimorphic, since the cells that differentiate as PVPL and PVPR in hermaphrodites instead become the PVU and PVS neurons in males, respectively (White *et al*. 1986; Cook *et al*. 2019). We found that in hermaphrodite PVP neurons—but not in males PVU and PVS neurons—the main neural processes sprout branches at the midbody that typically extend dorsally and terminate in large wing-like structures that lie over the connection between the uterus and vulva. This is the connection through which hermaphrodites receive sperm into the uterus when they are mated by males, and through which eggs pass out of the uterus when they are laid. The PVP branches are hermaphrodite-specific since they are not seen in the male PVU or PVS neurons. Further, the PVP branches develop at a time during the L4 stage when other sexually dimorphic structures of the hermaphrodite also develop, such as the vulva and the neurons of the egg-laying circuit (Desai et al. 1988; Mok *et al*. 2015). These observations are all consistent with the hypothesis that the PVP neurons carry out functions related to adult hermaphrodite reproduction, such as mating or egg laying.

### Are the PVP neurons ciliated sensory neurons?

The PVP branches terminate in structures that could be sensory cilia, although the evidence for this remains equivocal. The wing-like termini of the PVP branches are about 10 μm across, and the most similar neural structures in the *C. elegans* nervous system are the “winged” cilia of the AWC amphid neurons, which occur at the termini of AWC dendrites near the nose of the animal. AWC neurons use these winged cilia primarily to respond to volatile odorants (Bargmann *et al*. 1993). Thus, we hypothesized that the PVP branch termini might also be sensory cilia. In favor of this hypothesis are three lines of evidence. First, the PVP branch endings morphologically resemble AWC cilia. Second, we found that the PVP branches and wings are surrounded by cells that express the *mir-228p::gfp* marker, which is commonly used as a glial-specific marker in *C. elegans*, and dendrites and cilia of known sensory neurons are similarly surrounded by glial cells that express this same marker (Shaham 2006). Third, the PVP neurons were found by Vidal et al. (2018) to express the sensory receptor homologs *srab-12* and *sri-9*. Interestingly, *sri-9* appears to be expressed in only dauer animals and then only in the left but not right PVP. This asymmetry between PVPL and PVPR is reminiscent of the asymmetries between the left and right members of *C. elegans* sensory neuron pairs such as ASE (Yu *et al*. 1997), PLM (Wicks and Rankin 1995), and AWC (Troemel *et al*. 1999), the latter of which are the sensory neurons with cilia similar in shape to the PVP wing structures.

Standing against the three lines of evidence that favor the hypothesis that PVP are ciliated sensory neurons are two types of experiments that gave negative results. First, the *daf-19* mutation, which disrupts the structure of cilia in known sensory neurons (Swoboda *et al*. 2000), had a far less striking effect on PVW. While ∼60% of the winged cilia of AWC neurons were disrupted in the *daf-19* mutant background, only ∼7% of PVW branches and wings were affected in the *daf-19* background. Second, we were unable to see GFP labeling of the PVW winged endings using transgenes that label sensory cilia in young larvae, although we were also not able to detect labeling of any of these sensory cilia with these transgenes in adult hermaphrodites, the only types of animals that have PVW branches and wings.

We conclude that it is a reasonable hypothesis that the PVP branches and wings are sensory dendrites and cilia, and that the surrounding structural cells of the egg-laying system may play a secondary role as glial cells that support these PVP structures. However, given that these would be quite surprising results to come up only now after decades of previous studies of the *C. elegans* egg-laying system, further experimental evidence is needed before a final conclusion can be reached about this issue.

### Challenges to studying PVP function

While we note that more experimental evidence about PVP function is required to make firm conclusions about its function, this article lacks such evidence because of technical challenges specific to the study of PVP. We tried multiple different promoters to express multiple different proteins in PVP, including fluorescent proteins, and we repeatedly found that these transgenes either failed to produce viable transgenic animals (perhaps due to disrupting the ventral cord pioneer function of PVPR), or resulted in viable transgenic lines in which PVP neurons had variable and “sick” appearing morphology. As a result, we determined that we could not reliably interpret results of experiments in which transgenes were used to manipulate or record PVP activity, which are the experiments needed to definitively determine the function of PVP. Our conclusions about the normal morphology of PVP are based on what we see most often in animals where PVP cells have a healthy appearance. Future progress in studying PVP function will require finding ways to manipulate and visualize PVP activity without making these neurons sick. Our efforts exclusively used multicopy transgenes, and perhaps single-copy transgenes or CRISPR engineering of endogenous genes will be more successful.

### What are plausible functions for PVP?

We did not detect any defects in egg-laying behavior in our transgenic animals carrying transgenes designed to inactivate PVP, but we did defect locomotion defects (data not shown). PVP is known to release the neuropeptide PDF-1 to promote long bouts of forward locomotion known as roaming, which contrast to an alternate pattern of locomotion in which animals make more reversals and stay in place, known as dwelling (Flavell *et al*. 2013). If PVP is a sensory neuron with a cilium, then it could sense mechanical or chemical signals from the hermaphrodite reproductive system over which the winged cilium lies. Therefore, PVP could sense something about mating or egg laying to regulate PDF-1 release and the switch between roaming and dwelling locomotion.

If PVP is a sensory neuron, one possibility is that it mechanically senses either eggs in the uterus, or the passage of eggs out of the uterus when eggs are laid. The accumulation of eggs in the uterus is sensed by a mechanism that remains unidentified (Collins et al. 2016), and the PVP could play a role in this mechanism. Egg-laying and locomotion behavior are coordinated by mechanisms that also remain undiscovered (Collins and Koelle, 2013), and again PVP could play a role in this process. Alternatively, as sensory neurons, PVPs could allow hermaphrodites to sense when they are being mated by males. For instance, the PVP wing structure could mechanically sense physical distortion from mating and contribute to regulating responses such evading males and/or expelling sperm (Kleemann and Basolo 2007). Since PVPs express chemoreceptors, it could also be the case that the PVP wing structures participate in pheromone signaling as part of the interplay between aging and mating (Ludewig *et al*. 2019; Aprison and Ruvinsky 2016; Maures *et al*. 2014). While these ideas remain hypothetical, they can direct further studies aimed at uncovering PVP function.

## Acknowledgements

We thank Dr. Richard Ikegami for supplying us with pEXP294, which contained the PVP-specific promoter referred to in this manuscript as PVPp. We thank the lab of Daniel Colón-Ramos for the *odr-1p::RFP* plasmid. This work was funded by National Institutes of Health Grant NS036918 to M.R.K. and a National Institutes of Health predoctoral fellowship F31NS115315 to N.T.M.C.

